# Modelling dopaminergic signals associated with habit formation through temporal-difference action learning

**DOI:** 10.64898/2026.08.10.743861

**Authors:** Charlotte Collingwood, Francesca Greenstreet, Marcus Stephenson-Jones, Rafal Bogacz

## Abstract

Action-selection is determined by a combination of goal-directed and habitual processes. Habits are defined as the reward-independent, stimulus-response relationships which form when an action is regularly executed in the same context, regardless of outcome. An influential computational model proposes that habit formation is driven by action prediction errors which occur when non-habitual actions are taken. It has been further suggested that action prediction errors are encoded in activity of specific dopamine neurons, and it has been recently observed that dopamine activity in the tail of the striatum follows a pattern consistent with the action prediction errors. However, the original models capture changes in habits across trials, but do not describe the time-course of action prediction errors within trials, hence it is difficult to directly compare them with dopamine activity. We begin by outlining the ‘temporal-difference action learning’ algorithm, which uses biologically-plausible mechanisms to determine how dynamic changes in action intensity influence the resultant prediction errors across near-continuous time. We then demonstrate that dopaminergic data recently collected from the tail of the striatum is better represented by action prediction errors than reward prediction errors. Overall, our results support the existence of value-free action prediction errors and associated habitual behaviour in dopaminergic signals.

**Author summary:** Whenever we choose one action over another, there are two ways that the selection can be made. We could take the time to consider what we want to achieve, calculate which action is the most likely to give us that outcome and balance it against the possible negative consequences. These ‘goal-directed’ calculations are very time-consuming and our brains could not possibly do it for every choice. Instead, we often rely on the second method, ‘habits’, which learn to copy the actions that were most often chosen in the past. In this paper, we present a new model of learning that is based on biologically plausible brain networks and applies *action* prediction errors to update our ‘habits’ across continuous time. Using simulations, we reveal testable predictions that are specific to our ‘temporal-difference action learning’ model and build an intuition for its behaviour. Finally, this model is tested against real dopaminergic data from the tail of the striatum, and we show that our model provides better explanation for these data, than classic ‘reward-based’ reinforcement learning models.

## Introduction

Colloquially, we have all experienced habitual behaviours and they are often blamed for the unintended actions we subconsciously make. A classic example is opening the fridge after walking into the kitchen, despite not deliberately looking for food. While this is a common scenario in the real world, it can appear strange on reflection - a sequence of actions are executed without any conscious control. What causes such *slips-in-action* in our behaviour when we aren’t paying attention? This is the question posed by researchers across many fields, including behavioural psychology, neurobiology and computational neuroscience, when they interrogate the nature of *habits*.

Maladaptive habits are associated with a wide range of medical disorders, either forming a core part of the root aetiology or as side effects from medical treatment. For example, habits are linked to compulsive drug seeking behaviours [1], gambling addictions [2], and the impulse control disorders associated with levodopa treatment for Parkinson’s disease [3, 4]. Our internal states are also strongly affected by habits - excessive rumination in depressed patients is often conceptualised as a ‘mental habit’ and modelled using a habit-goal framework [5]. Similar arguments are given for obsessive compulsive disorders [6] and motor compulsivity in Tourette’s syndrome [7]. Striatal dopamine has been particularly implicated in all of the above disorders and has long been thought to play a role in habit formation [2, 4, 6–9]. This is partially due to the historic association of the basal ganglia (BG) as a brain region which both (1) controls reward-learning through dopamine [10] and (2) is responsible for the expression of goal-directed and habitual actions [11]. Aligning the evidence of dopamine’s role in habit formation with the mathematical models describing this process is likely to play a key role in developing future treatments for disorders in which harmful habits form and become compulsive.

A key theory for how one action may be chosen over another proposes that the filtering of potential action plans is done through a two-process system [12]. The *goal-directed* system takes time and resources in order to analyse the current context and the likely outcomes of all available choices, then makes the choice that either maximises the positive consequences or minimises the negative. Though this goal-directed system is flexible and effective, the brain has thousands of choices to make every second and insufficient resources to calculate the goals and outcomes for all of them. This is where the habit process is useful, as it learns which choice is usually made in a given context and copies previous actions, thus allowing for rapid decisions to be made with minimal analysis.

Until recently, the exact definition, and consequent modelling, of what constitutes a habit has been inconsistent between the experimental and computational neuroscience fields. In psychological and behavioural studies, where unconscious slips-in-action were first established, habits are defined as the reward-insensitive stimulus-response (S-R) behaviours [13]. These develop in extended instrumental conditioning experiments [14], during which animals repeatedly perform the same action in response to a stimulus until this behaviour becomes automatic. This automaticity eventually leads to inflexibility - when a choice is no longer optimal, habitual actions will take much longer to extinguish than those which are still goal-directed [14]. In contrast to S-R theory, the computational neuroscience field often represents habits with model-free reinforcement learning (RL) algorithms (as the alternative to model-based goal-directed choices) [15]. The model-free system is assumed to learn optimal action policy by minimising the reward prediction error (RPE), and it selects choices with the maximum expected value in response to a given stimuli. However, a model-free system that *relies* on RPE is fundamentally incompatible with the reward-insensitive nature of habitual S-R behaviours.

This discrepancy in definitions was addressed by Miller et al. [16], who proposed that this computational ‘model-based or model-free’ understanding of action-selection processes should be replaced by a ‘value-based’ versus ‘value-free’ dichotomy. Their argument was two-fold. First, they asserted that the evidence for dissociable neural correlates for model-based and model-free signals is inadequate [17–19], especially in comparison to the dorsal striatum (DS) lesion experiments which cleanly separated goal-directed actions from habits [11, 20–22]. Second, they restated an argument previously given by Dezfouli and Balleine [23] - model-free RL algorithms cannot reproduce the experimental results of resistance to action-outcome (A-O) contingency degradation tests. Put simply, the model-free system learns the action-value of a given state. Therefore, it *should* have the capacity to learn that execution of an action now leads to a negative outcome (reward delay) and that selecting to perform ‘no action’ is more valuable. This suggests that the previous model-based/model-free dichotomy is insufficient to describe the separation of habits and goal-directed behaviour.

Miller et al. [16] argue that the answer to describing habit formation lies in replacing RPEs with *action* prediction errors (APEs). Conceptually, APEs provide an intuitively appealing solution for an S-R learning mechanism, where S-R habits learn to mimic actions which have previously been executed in the current context. So, by its nature, the prediction error (PE) in such a system must measure how successfully the agent expected the antecedent action, rather than the outcome. For example, in an instrumental association task, the first time an action is made in response to a cue, a large APE would occur and the habit system would update to expect that particular action slightly more when the cue is next experienced.

The precise formulation for an APE-based habit system is not fixed, but some properties are fundamental. An APE should (1) display a peak following unexpected actions, and (2) be entirely reward-insensitive. Originally, two key models have been developed that employ APEs. Namely, Miller et al.’s [16] value-free habit system and Bogacz’s [24] DopAct model, which additionally proposes that APE signals are encoded in the activity of a subset of dopamine neurons that drive habit formation. Both models are able to replicate reversal learning, devaluation and contingency degradation effects, while explaining the influence of different schedule of reinforcements (SORs) on habit strength [25]. However, these two proposed models worked on a trial-by-trial basis and neither developed a mechanistic model for how such APEs could be produced and evolve within a trial, nor directly related them to experimental data on the dopamine dynamics in the BG.

This paper extends upon the models proposed by Miller et al. [16] and Bogacz [24] to address two questions:

1. Can habit learning be generalised to continuous and scalar actions?
2. Can evidence of action prediction errors be found in striatal dopamine?

To answer these questions, this paper presents the temporal-difference action learning (TD-AL) algorithm, a biologically-plausible learning model which produces predictions of expected (future) actions, based on prior experience. Further, TD-AL’s performance is tested and compared to striking neural evidence of APEs recently reported by Greenstreet et al. [26].

Rather than learning total expected reward, TD-AL predicts the total expected (future) action intensity, and learns it using an APE - a prediction error signal analogous to RPEs but based on action rather than reward. These actions intensities can be continuous; the model is able to predict *when* the action will occur, relative to predictive cues, and it produces realistic APEs which abide by standard theories of dopamine signalling. Its algorithm can also be extended to explain a variety of behavioural and neural data. In doing so, this work demonstrates how one principle, namely TD learning, can capture learning in diverse striatal regions.

An earlier version of this paper was presented in an abstract form in the proceedings of Computational Cognitive Neuroscience conference [27].

## Results

This section is structured into three components. It begins with the mathematical definition of TD-AL together with a description of the underlying conceptual framework. Then, an intuition for the learning dynamics is built through the simulation of a classic instrumental association task. Finally, TD-AL is compared to the established temporal-difference reinforcement learning (TD-RL) model in their capacity to model and reproduce real neural data.

### Model description

TD-AL is based on Sutton and Barto’s [28, 29] TD(*λ*)-RL model. This section outlines the mathematical adaptations that are implemented to introduce realistic neural representations of an APE and the theoretical assumptions that motivated these choices.

#### Theoretical assumptions

Given the parallel connectivity and structure of the striatum [30, 31], it has often been posited that these loops represent a ‘computational unit’ since they should, logically, manipulate the information they are given in an identical manner [24, 32–34]. TD-AL follows this formulation and the assumption that the information being predicted in any given region of the striatum is determined entirely by the outcome that influences the corresponding prediction error signal, which may vary along a gradient from pure reward (e.g., in the ventral striatum (VS)) to pure action (e.g., the dorsolateral striatum (DLS) or, perhaps, the tail of the striatum (TS)).

This further requires the foundational assumption that dopamine performs an identical computational role throughout the BG. More precisely, to maintain a high degree of biological plausibility, TD-AL argues that:

1. Phasic dopaminergic dynamics encode a neural prediction error signal.
2. The dopaminergic signal determines the direction and magnitude of synaptic plasticity changes in the striatum.
3. The prediction errors can be produced by disparate events in the environment.
4. The BG is formed of parallel computational units and the information in any given system is determined by its input.

Consideration of how an action is defined is particularly relevant in the context of habits, as maladaptive expression is rarely associated with the execution of a single key-press. For the purposes of the work presented here, choices and action plans are treated as synonymous, each with it’s own *H_a_*; for example, classifying a choice as pressing the ‘left’ or ‘right’ lever, rather than total relative position or individual changes in muscle groups. For an action in near-continuous time, this is represented by a single continuous curve which ramps up and down as movement is initiated and ended.

This simplification is based on, and validated by, prior work into goal-directed action modelling [16, 24, 35, 36] which states that the BG are responsible for filtering between plans presented by the cortex, rather than controlling the mechanical execution of the action - a process which can occur downstream and is more often attributed to the cerebellum [37, 38] (another site of much neuroscientific RL research [39–41]).

TD-AL learns from action *intensity* with the implication that the striatum is, to some degree, responsible for the *vigour* with which an action plan is executed. This aligns with the historical studies associating striatal dopamine with motivation and execution of planned movement [42–46], but requires the striatum to not only act as a ‘gate’ for action initiation but also to take into account factors other than just the intended movement and outcome.

#### Learning to predict actions

As in previous models [16, 24], we assume that habits are learnt similarly to values in TD-RL. To represent an action-based value-free habit, TD-AL learns to estimate the action intensity, *A_a_*(*t*), by minimising prediction errors. The optimal state-value of *H_a_* is denoted by *H*_*a*_^*^ and is calculated using Eq 1. Specifically, for each potential action, *a*, the ‘striatal’ activity predicts future expected action intensity, *H_a_*, given the current state. *A_a_* is continuous and can last (i.e., have a non-zero value) for several timesteps.

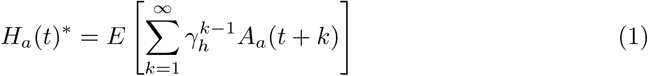

where: *H_a_*= expected future intensity of action, *a*,

*γ_h_* = action discount factor,

*A_a_*(*t*) = action intensity of action, *a*, at timestep, *t*.

Consequently, the computational structure of the prediction error itself is unchanged from the TD(*λ*)-RL RPE, the outcome variable is simply altered from reward to action intensity. At each timestep, the APE is produced:

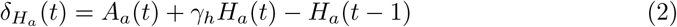

where: *δ_Ha_* = action prediction error.

#### Credit assignment and microstimuli

A critical new feature of this model, relative to previous APE formulations, is its capacity to predict *when* an action usually occurs after the associated stimulus has been perceived. The strength and behaviour of habits are time-dependent [13] and variable SORs are more likely to result in delayed extinction behaviour [47]. Therefore, the ability to interrogate how temporal delays influence the strength of habit versus goal-directed actions is advantageous.

To achieve this, TD-AL utilises the ‘microstimulus’ representation of an eligibility trace, as proposed by Ludvig et al. [48, 49], to model the expected interval between cue and action. This algorithm was selected for its high biological-realism and improved replication of complex dopaminergic dynamics, relative to the standard decaying *λ* memory trace [48]. This formulation has been increasingly applied over the last decade [50–53] and, unless otherwise stated, all future mentions of ‘TD-RL’ in this paper will rely on microstimuli representation.

The microstimulus algorithm states that, following the occurrence of a stimulus, a series of *m* microstimuli is produced, which are each represented by Gaussian basis functions. In combination, these microstimuli track the decaying stimulus trace over time by providing overlapping ‘temporal receptive fields’.

Mathematically, a decaying memory trace, *y_i_*, is produced in response to each salient event in the environment (Eq 3). This is similar to the standard eligibility trace in TD(*λ*)-RL model [28, 29, 54]. This trace is convolved with a uniformly distributed sequence of Gaussian curves (Eq 4) which results in the final microstimuli (Eq 5).

Note that this algorithm produces curves with uniformly distributed peaks (as *µ* = *j/m*) and increasing widths, which replicates the decreasing temporal accuracy as the delay increases seen in striatal neurons [55],

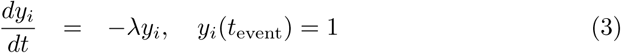

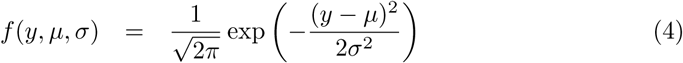

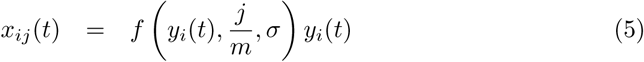

where: *λ* = memory decay parameter,

*y* = memory trace,

*t* = timestep,

*i* = stimulus,

*µ* = Gaussian centre,

*σ* = width of each curve,

*j* = microstimulus,

*m* = total number of microstimuli for any stimulus, *i*.

Thus, for *every* event, *i*, that occurs (e.g., cues, actions and rewards), *m* microstimuli are produced. The total habit value, *H_a_*, at any given time is composed of the current microstimulus activity, *x_ij_*, modulated by their relative weights, *w_H_a_,ij_* (Eq 6). The weights are then updated proportionally to both the prediction error produced and the activity of the associated microstimulus (Eq 7). The equivalent equations for a TD-RL system are adapted from Ludvig et al. [48] and are given below (Eq 8 and Eq 9).

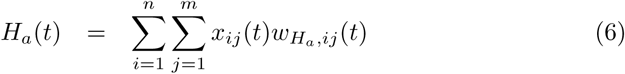

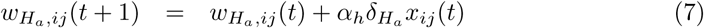

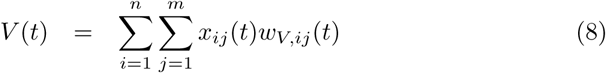

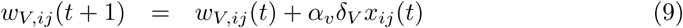

where: *w_H_a_,ij_* = *H_a_*weight for the *j*^th^ microstimulus caused by event, *i*,

*α_h_* = habit learning rate,

*n* = total number of events experienced,

*w_V,ij_* = *V* weight for the *j*^th^ microstimulus caused by event, *i*,

*α_v_* = value learning rate.

Note that the microstimuli activity levels, *x_ij_*, are independent of the learning model as they represent the external state/context. For a heterogeneous BG, the same microstimuli would be shared across all habit and value variables. It is the weights that contain the information learnt about the measured outcome (i.e., the specific action/reward). All microstimulus parameters are fixed to the values listed in Table 1. Predictive cues provide microstimuli with steadily increasing weights which causes the model’s expectation of future action intensity to increase. Note that some action microstimuli are negatively weighted, as they counterbalance the positive weighting of the other microstimuli to return *H_a_* to 0.

**Table 1.**
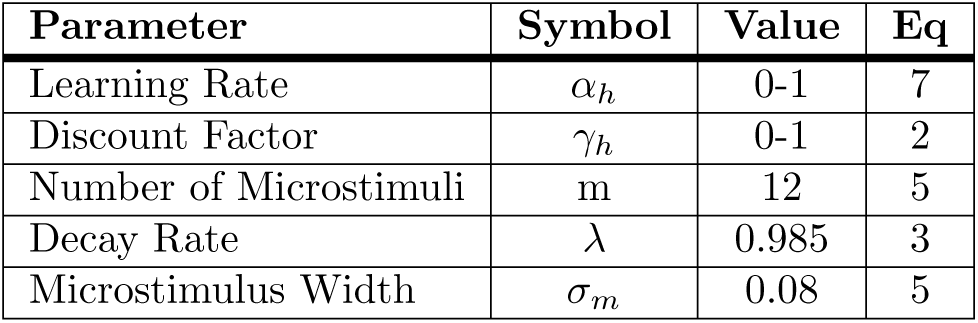
TD-AL parameter values, including microstimuli.

| Parameter | Symbol | Value | Eq |
| --- | --- | --- | --- |
| Learning Rate | $\alpha_h$ | 0-1 | 7 |
| Discount Factor | $\gamma_h$ | 0-1 | 2 |
| Number of Microstimuli | m | 12 | 5 |
| Decay Rate | $\lambda$ | 0.985 | 3 |
| Microstimulus Width | $\sigma_m$ | 0.08 | 5 |

In their initial formulation, Ludvig et al. [48, 49] included an eligibility trace for each *x_ij_*. However, we found that the impact of including this was minimal given that *x_ij_* is already spread across time and is sufficient to resolve the credit assignment problem for our purposes. Consequently, it was not included in this paper’s algorithms.

Fig 1A and Fig 1B demonstrate the dynamics of the microstimuli produced for a single ‘cue’ event and a cue-action-reward series, respectively. Fig 1B helps to provide an intuition for what Eq 6 computes. Each microstimulus’ weight, and thus, its contribution to the final action expectation, is represented by its relative opacity. Their weighted sum results in the ‘habit’ curve shown.

**Fig 1.**
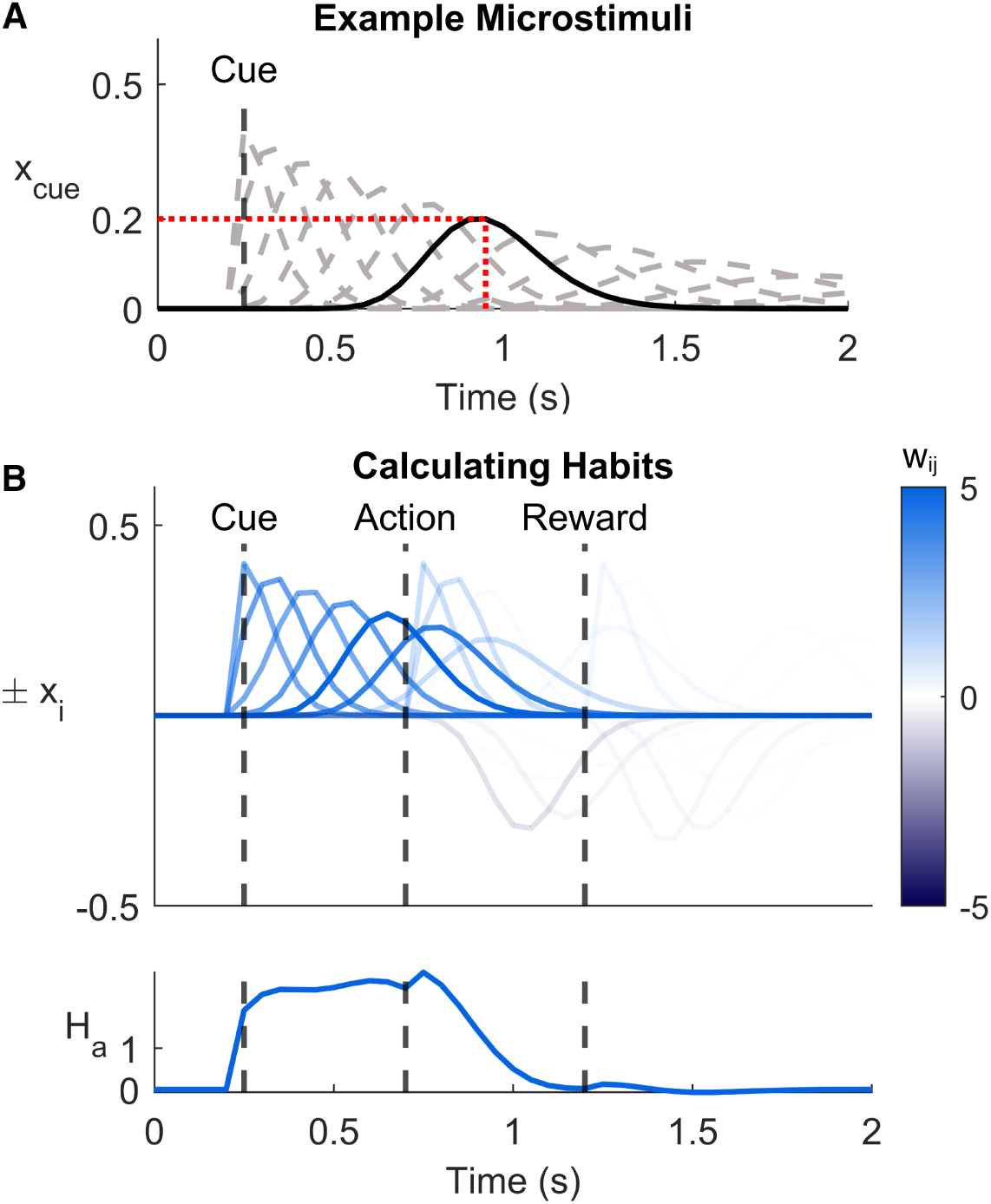
Representation of expected action through microstimuli. **A**: An example of microstimuli dynamics following a single ‘cue’ event. Parameters are taken from Table 1. The peak firing rate is uniformly distributed and decays across time. The activity of microstimulus *x*_cue,7_ (solid line) at 0.95 seconds after the event is 0.2, as demonstrated by the red line. **B:** Together these two panels portrays how the habit function is produced by the microstimuli. (Top) An example simulation of a single trial, with three events: cue, action and reward. Each event produces its own microstimuli. The opacity is adjusted to their relative weighting after TD-AL is trained on 200 trials (*α_h_* = 0.2*, γ_h_* = 0.97). Microstimuli with negative weights are plotted below the zero line for additional clarity. (Bottom) The *H_a_* function resulting from the same simulation.

The microstimulus model confers several advantages. Primarily, it aligns with our current understanding of striatal connectivity and is more mechanistic than the standard eligibility trace. For example, the microstimuli could be implemented by cortical inputs to the striatum or striatal sub-populations with different temporal dynamics following salient stimuli. Such time-dependent firing patterns have been observed in the BG’s medium spiny neurons [55, 56] and cortical oscillators have previously been proposed as the source of a striatal ‘internal clock’ [57, 58]. Further, dopamine influences the strength of synaptic connectivity via Hebbian learning, which can be concretely correlated to these individual weights [59–61].

### Simulations of instrumental association learning

This section helps to build an intuition for how TD-AL’s algorithm behaves through the simulation of a classic experiment in which the activity of dopamine neurons is often recorded [10, 62].

Instrumental association experiments have been key in developing TD-RL and S-R theory [13, 63, 64]. In their simplest form, an animal learns to perform an action in response to a neutral stimulus to gain a reward by repeatedly experiencing a cue-action-reward sequence (as in Fig 1B) and developing an estimation of both S-R and A-O contiguities [13, 65, 66].

Three models, whose key characteristics are outlined in Table 2, are compared to interrogate two factors:

1. How does the APE differ from the better-known RPE (i.e., TD-RL vs. TD-AL*_γ_*)?
2. What is the influence of the *γ_h_* term on both the APE and *H_a_* variable (i.e., TD-AL*_γ_* vs. TD-AL_0_)?

**Table 2.** Key characteristics of the three simulated models.

| Label | Algorithm | Outcome | $\alpha$ | $\gamma$ |
| --- | --- | --- | --- | --- |
| TD-RL | TD-RL | Reward | $\alpha_v = 0.2$ | $\gamma_v = 0.97$ |
| TD-AL$_{\gamma}$ | TD-AL | Action intensity | $\alpha_h = 0.2$ | $\gamma_h = 0.97$ |
| TD-AL$_0$ | TD-AL | Action intensity | $\alpha_h = 0.025$ | $\gamma_h = 0$ |

Note that $\alpha_h$ has been decreased for TD-AL $_0$ to keep the size of weight updates easily comparable between models.

In the following simulations, all three models experienced 200 instances of the cue-action-reward sequence, with a delay of 10dt between each event (1dt = 0.05s) and a 7.5s inter-trial interval (ITI). The results of their learning are shown in Fig 2.

**Fig 2.**
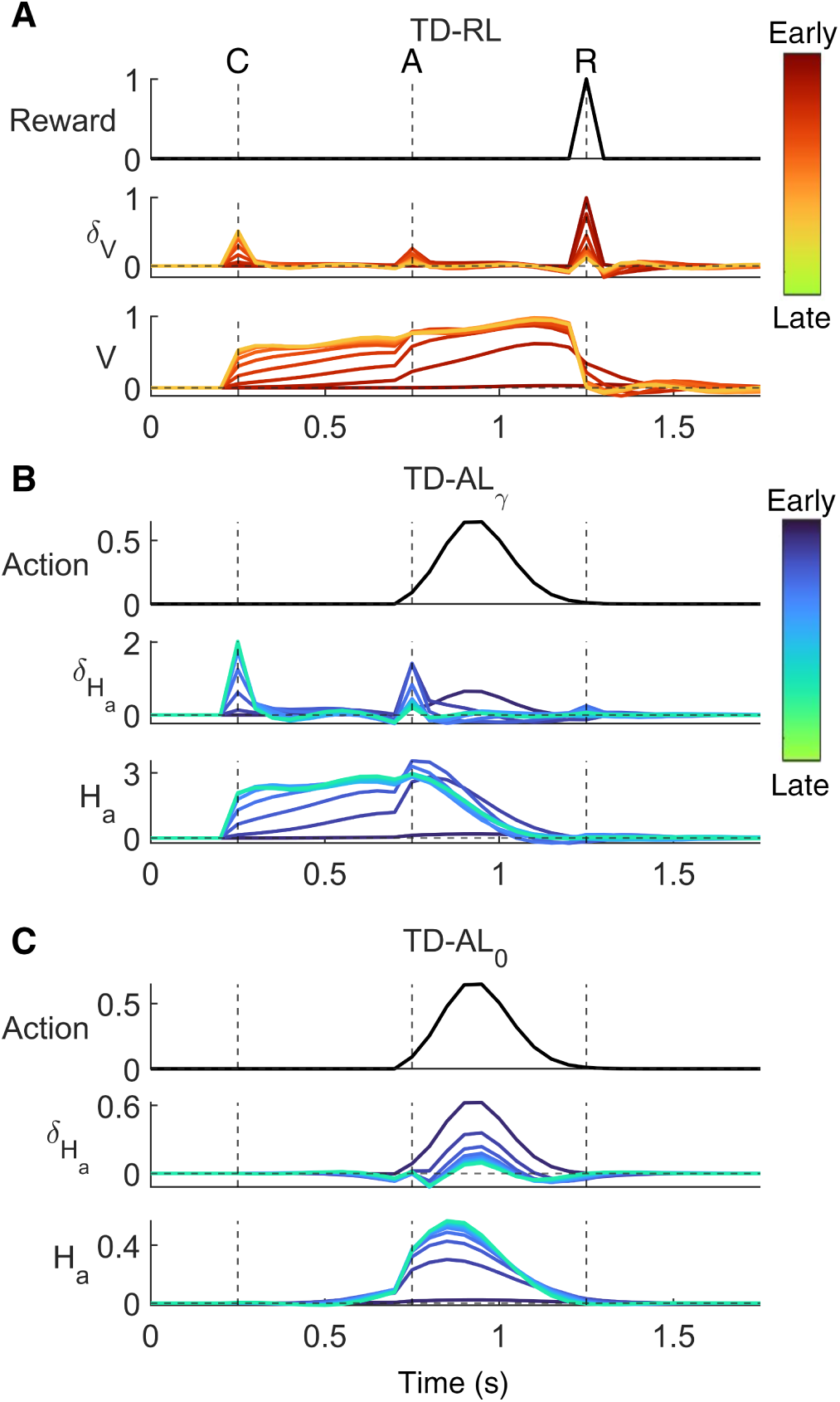
Instrumental Association Simulations. The evolution of three models learning a simulated instrumental association task. Each model experienced 200 trials, where cue, action initiation and reward were 0.5s apart, with a 7.5s ITI. (Top) The measured outcome value. (Middle) The prediction error from early to late trials. (Bottom) The estimated outcome learnt by the model from early to late trials. **A:** The results of a TD-RL model (*α_v_* = 0.2*, γ_v_* = 0.97) from early trials (red) to late in learning (yellow). **B:** The results of a TD-AL*_γ_* model which learns to predict upcoming continuous action intensity (*α_h_* = 0.2*, γ_h_* = 0.97) from early trials (blue) to late in learning (green). Both *δ_Ha_* and *H_a_* are insensitive to reward. **C:** As TD-AL*_γ_*, but learning instantaneous action intensity rather than future action (*α_h_* = 0.025,*γ_h_* = 0).

#### TD-RL vs. TD-AL***_γ_***

TD-RL behaves exactly as predicted of a reward-based reinforcement learning model. With training, the RPE transfers from unexpected reward to earlier predictive cues before converging on the earliest and *V* learns to increase with each consecutive cue until the expected reward is received. The temporary response to action initiation results from the explicit assumption that the action itself triggers its own set of microstimuli and therefore behaves as a predictive cue.

In contrast, TD-AL*_γ_* shows all the features predicted for an APE: (1) it is movement-locked to the initiation of the unexpected action, and (2) it is reward-insensitive. Additionally the APE shifts to the predictive cue as learning occurs. Due to the continuous Gaussian action profile, *δ_Ha_* initially copies the dynamic structure of the action intensity itself (black curves in Fig 2B) until *H_a_*’s predictions improve and are better able to approximate ‘future remaining action intensity’.

#### Effects of the discount factor

In TD-RL, *γ_v_* determines the degree to which the model expects or discounts future rewards. However, when considering actions, this parameter can arguably be considered superfluous. It is unclear whether it would be more beneficial for the striatum to predict which action should be made *right now*, or to prepare for the amount of action, and thus future effort, that is usually executed in its current context.

Given that a key theoretical proposal for the function of the basal ganglia includes a VS-DLS gradient of representations, from reward through goal-directed selection to habits [67] across a common computational unit, TD-AL makes no claim as to the likely value of *γ_h_*. The degree to which future outcomes are discounted could vary across striatal regions, and so, the inclusion or exclusion of *γ_h_* are not incompatible outcomes - both versions of the learning algorithm could be utilised in the striatum to different ends. Making a prior assumption of *γ_h_* will only be necessary if *H_a_* is used to predict an animal’s actions, as it influences our interpretation of the variable.

Comparing TD-AL*_γ_* and TD-AL_0_ allows us to explore how a *γ_h_* value of 0.97 or 0 will influence the evolution and final structure of *H_a_* and *δ_H_a__*. Two key features develop. First, as shown in Fig 2B, *H_a_* behaves similarly to *V* in TD-RL for TD-AL*_γ_*. Action expectation increases when the predictive cue occurs and the expected amount of future action intensity is calculated once the action has begun. TD-AL*_γ_* demonstrates this with a gradual decrease in *H_a_* after action initiation, which integrates to the total amount of movement that has not yet occurred, rather than the sharp drop visible for TD-RL. In contrast, TD-AL_0_ learns to predict in *H_a_* the immediate continuous structure of the action (here, a Gaussian curve), as well as it can be approximated from a Gaussian microstimulus basis function.

Second, TD-AL_0_’s *δ_Ha_* does not show the transfer of dopamine prediction errors to the earliest predictive cue - despite this being a key characteristic demonstrated in TD-RL. Instead, the APE converges to 0. However, this does not mean that experiencing the cue has no influence on action expectation, as proven by the increase in *H_a_* just prior to initiation. Even with *γ_h_* set to 0, the cue microstimuli have non-zero weights and the presence of these microstimuli causes an expectation of action to build. As such, the cue is still vital to the prediction of action timing, but the microstimuli are not linked to an instantaneous action change, and so, no PE occurs.

Intriguingly, this formulation therefore implies that, when an S-R relationship has been learnt, TD-AL_0_ predicts that no associated dopaminergic signals would be present or detectable. As such, cue-locked dopamine activity is no longer a requirement to determine that prediction-error-driven learning is occurring.

### Modelling neural data

This section provides both (1) a proof-of-concept that TD-AL can be used to test if real neural data encode APEs and (2) evidence that supports the existence of APE signals in rodent dopaminergic recordings. To begin, a description of the experiment and relevant results from Greenstreet et al. [26] is provided, followed by an analysis of the extent to which these data can be explained by TD-AL or TD-RL, using Bayesian model selection (BMS) methods [68].

#### Experiment suggesting APE in the tail of striatum

Greenstreet et al. [26] recently published dLight recordings from the TS that appear to exhibit all the properties required of a dopaminergic APE signal, i.e., they are movement-locked, decay over trials and are unchanged by the presence or omission of a reward. Thus, their study presents a rich dataset with the potential to confirm the existence of APEs in the brain. A brief overview of the pertinent experimental methods and results is given below and schematised in Fig 3. A full description is provided in Greenstreet et al. [26].

**Fig 3.**
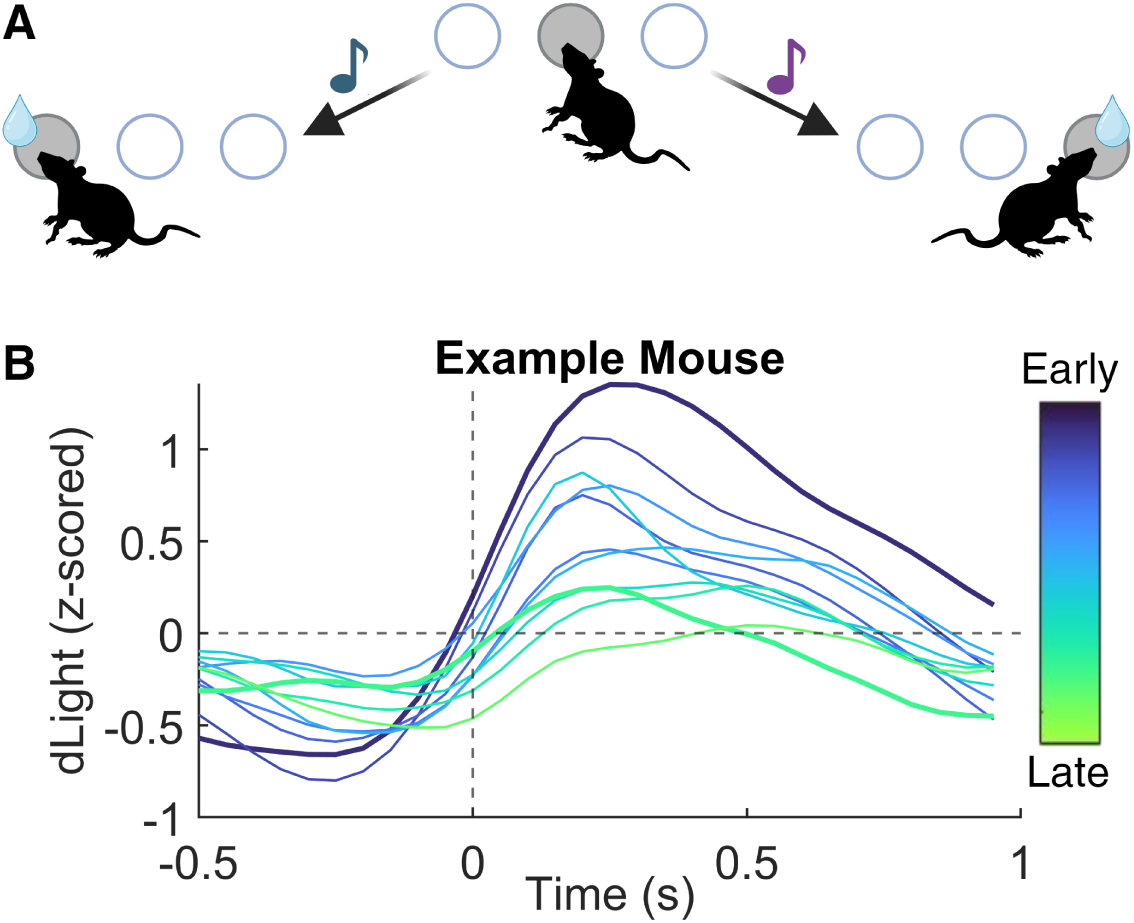
Experimental Design and Results of the study by Greenstreet et al. **A**: A schematic overview of the ‘cloud of tones’ task. Trials are self-initiated through a held nose-poke in the centre port (100-300ms), then an auditory cue indicates which port will produce a reward following a nose-poke. **B**: A recreation of the example mouse dLight recordings in the original paper, aligned to movement initiation. Each curve is an average of 200 consecutive trials, from early (blue) to late (green). The first and final 200 trials are indicated by thicker curves.

This dataset contains the dopaminergic signals of water-deprived mice in either the VS or TS during a two-choice ‘cloud of tones’ task [69] (Fig 3A). The mice were placed in a box with ports on three walls (left, centre and right) and could self-initiate a trial by holding a nose-poke in the centre port for 100-300ms whenever a centre LED was lit. A successful initiation poke resulted in the centre LED turning off and an auditory cue sounding 0-50ms later. The physical distance between the ports and the hold requirements were designed to increase the temporal separation between cues, actions and rewards. The frequency of the auditory cue (high or low, selected between two octaves) indicated which port (left or right) would produce a reward (2*µ*l of water) following the corresponding nose-poke. The mapping between frequency and port was counterbalanced between animals. Mice completed this task for over 10,000 trials and successfully learned to differentiate high- and low-pitched tones. For the training task that produced the data used in this paper, water-delivery was deterministic - as long as the mouse made the correct choice, it would receive a reward.

Greenstreet et al. [26] employed a fluorescence recording method to measure dopamine in which an artificial dopamine receptor, dLight, is injected into the relevant neural region using a viral vector [70]. These receptors emit a known amount of light for each molecule of dopamine that binds to them, allowing the quantity of the monoamine in the given area to be measured. Photometric dLight recordings began immediately after the initial habituation period. The researchers elected to record from two neural locations, the VS and TS. The former acted as a control and showed the clear RPE signals that have been regularly reported and are well-established [71].

Interestingly, despite the historical association of DLS with expression of S-R behaviours [11, 72, 73], this study chose to interrogate the TS, rather than the DLS. This decision was motivated by the fact that, of all the striatal structures, the TS is located the furthest away from the reward-associated VS and receives no efferent signals from the ventral tegmental area. As such, it is the least likely region to express RPE signals, which may otherwise mask the presence of APEs.

Their results aligned with the properties expected from an APE; TS dopamine expressed a peak that was locked to movement initiation, decreased across trials from early to late in learning (Fig 3B), and the TS signal was reward-insensitive.

Thus, the dopaminergic and striatal signals are action-sensitive, play a role in habit formation and demonstrate key features of an APE. We extend on this body of work by interrogating whether the complex dopaminergic dynamics can be explained by our mechanistic APE model.

#### Candidate models

The dopaminergic data was used to study two key factors:

1. Does TD-AL, rather than TD-RL, provide the best fit to the data?
2. Is there evidence for a temporal discount factor, *γ*, in the data?

In total, four model combinations were fit to the data, with a mouse’s speed being utilised as a proxy for action intensity. Their formulations and equations are outlined below and summarised in Table 3. The first three of these match the algorithms used for the instrumental association simulations above. The fourth model is a control capturing an alternative hypothesis that dopamine encodes movement speed and the collapse in dopamine over trials could simply be a reflection of the movement kinematics, which improve with training, rather than a true representation of a learning signal. Previous studies have detected dopaminergic responses to movement vigour and initiation [74–76] and performance on this task speeds with experience. Thus, it is not implausible that dopamine responses to movement would appear to collapse as the time to travel between ports decreases across learning. To account for this possible effect, the fourth model contains no learning algorithm at all. This *action-only* model is equivalent to setting the learning rate, *α*, to 0. As a result, the microstimulus weights, *w_ij_*, never update and the PE becomes a direct measure of the mouse’s speed (Eq 11).

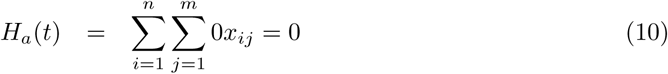

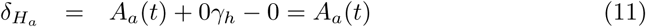

**Table 3.** An overview of the four models tested on Greenstreet et al. [26] data.

| Label | Algorithm | $\gamma$ | $\alpha$ | $\kappa$ |
| --- | --- | --- | --- | --- |
| TD-RL | TD-RL | Free (0-1) | Free (0-1) | analytic |
| TD-AL$_{\gamma}$ | TD-AL | Free (0-1) | Free (0-1) | analytic |
| TD-AL$_0$ | TD-AL | 0 | Free (0-1) | analytic |
| Action-only | TD-AL | 0 | 0 | analytic |

Note that a third parameter, $\kappa$ , is included. This is a scaling parameter and has no impact on learning behaviour (see Methods).

#### Model and parameter recovery

Before applying the fitting procedure and statistical analysis described in Methods to the real data, we tested the reliability of this analytical pipeline. As we do not include an algorithm through which *H_a_* is translated into future actions, we cannot produce fully simulated behavioural datasets. Thus, a more specific form of parameter and model recovery was undertaken by creating simulated PE signals from the behavioural data of each individual mouse across a systematic range of parameters, as outlined in Methods. These simulated data are created by ‘learning’ from the mouse’s experience, including cue timings, action intensities and rewards, using the same ‘behavioural simulation’ method as for the true fitting procedure. In doing so, they provide an indication of the degree to which the data available for each mouse are sufficient to constrain parameter values and select the correct model. The combined results for all mice datasets is provided in Fig 4.

**Fig 4.**
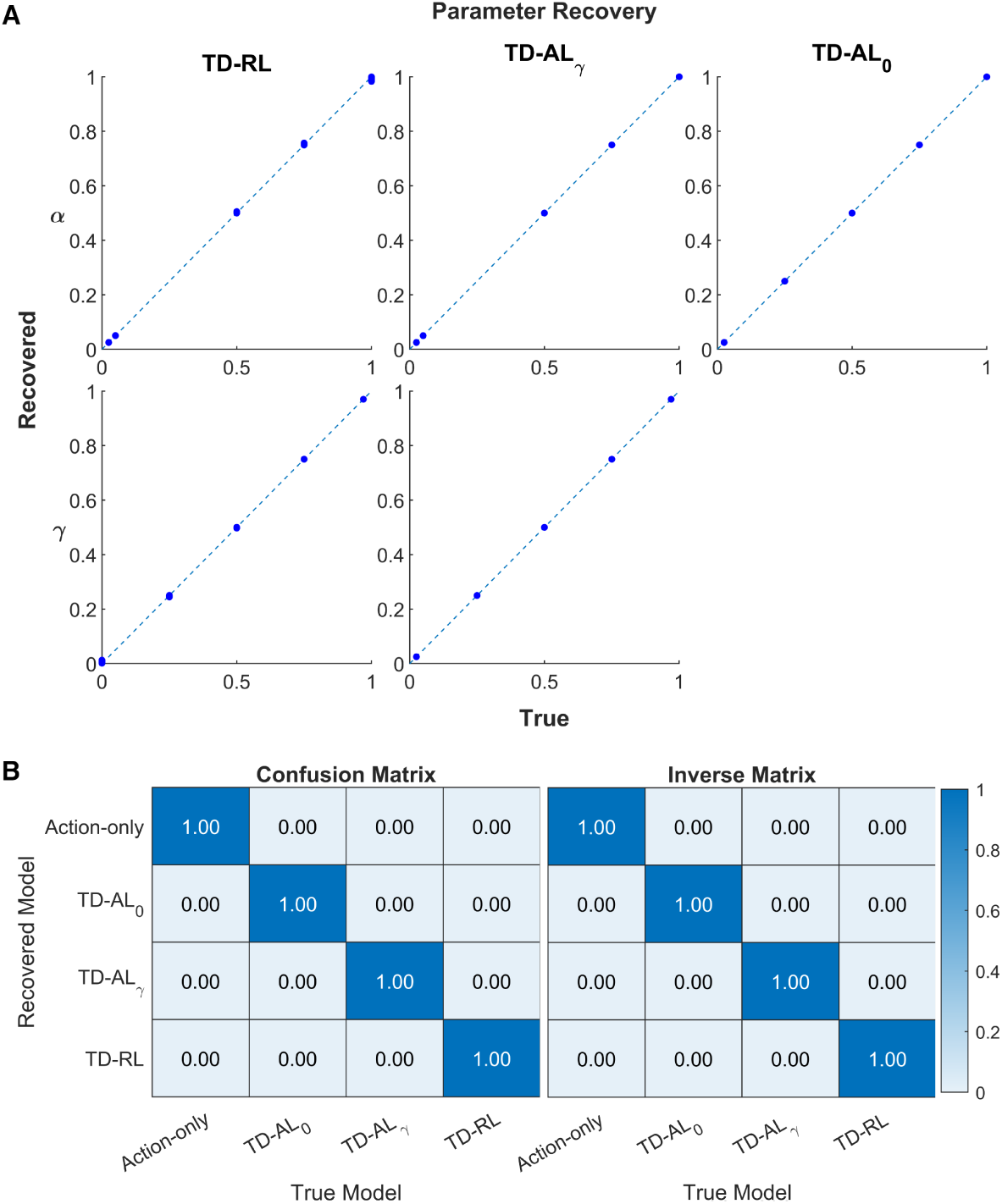
Parameter accuracy and model recovery matrices for simulated datasets. **A:** Parameter recovery analysis via the the correlation of recovered parameter value in relation to the true values used for simulations. From left to right: TD-RL, TD-AL*_γ_*, TD-AL_0_. All models have near-perfect linear correlation. **B:** Model recovery analysis. (Left) The confusion matrix, showing the proportion of models recovered (y-axis) given the true model (x-axis). (Right) The inverse matrix, showing the probability of the underlying model *given* the model that was recovered. All models were recovered correctly.

These show that the fitting procedure applied in this paper is consistent and likely to produce accurate results. The recovered parameters are linearly correlated with the true value used to simulate the data, for all mice, models and parameter combinations. The confusion and inverse matrix further reveal that all models could be distinguished from each other at the tested parameter combinations. We therefore conclude that this fitting procedure produces accurate measures of parameter values and underlying model.

#### Individual mice

The fitting procedure and statistical analysis were then applied to determine the best-fitting model for each individual mouse. The similarity of the true data to the prediction errors produced from each model’s best fitting parameters is illustrated (Fig 5) to allow the success of recovery to be assessed. Should none of the simulations provide qualitatively good results then the statistical Bayesian information criterion (BIC) analysis would simply reveal which model was ‘least poor’, rather than providing evidence for a given underlying mechanistic model.

**Fig 5.**
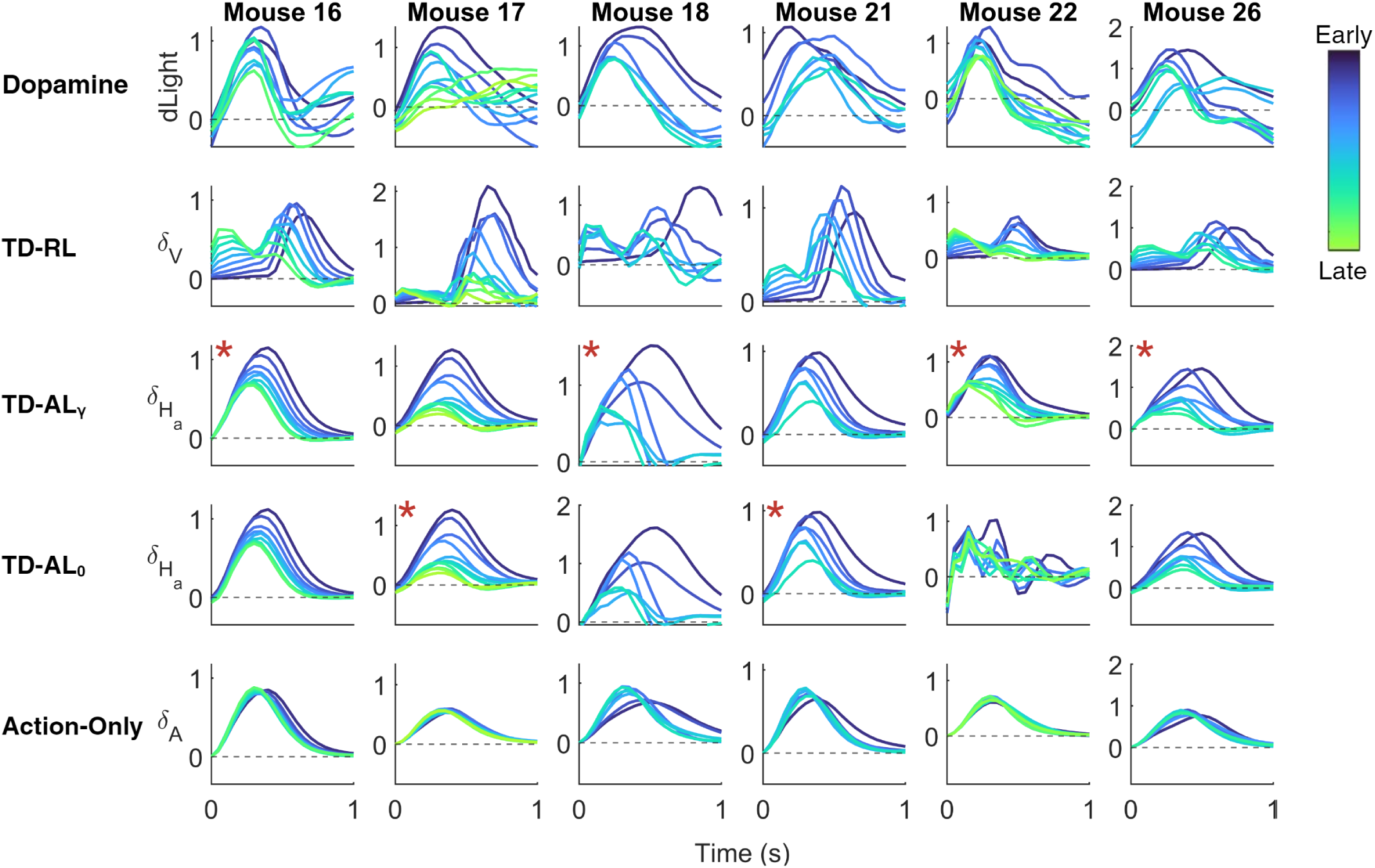
Comparison of dopamine activity with prediction errors from different model. Each panel represents the corresponding signal from early (blue) to late (green) in learning, aligned to action-initiation. Each line represents an average of 200 consecutive trials (there are a different number of curves for different animals because some mice had fewer trials with valid recordings than others). Each column contains the results for a given mouse, and the best-fitting model is indicated using a red asterisk, . The first row provides the true dopaminergic signal. The other rows show the simulated and convolved PEs produced by the best fitting parameters (Table 5). From top to bottom: TD-RL, TD-AL*_γ_*, TD-AL_0_, the action-only model.

Visual analysis confirms that TD-RL provides a poor fit for these data as the simulated results bear little to no resemblance with the dopaminergic signal. A significant delay between action-initiation and reward was enforced, so the corresponding RPE peaks arrive much later than is seen in the dopaminergic data. Further, if they contain any structure at all, the peaks sometimes increase in size across learning. In contrast, the two TD-AL models and the action-only model largely appear to qualitatively capture the timing of the peaks and the pattern of collapse. Therefore, it is appropriate to continue and interrogate the degree to which these models quantitatively replicate the data. The best fitting model and corresponding BIC and Akaike information criterion (AIC) results for each mouse are provided in Table 4.

**Table 4.** The BIC/AIC values and associated preferred model produced by the best-fitting parameters for each mouse. The lowest value for each mouse is highlighted in bold.

|  | Mouse | TD-RL | TD-AL<sub>γ</sub> | TD-AL<sub>0</sub> | Action-Only | Preferred Model |
| --- | --- | --- | --- | --- | --- | --- |
| BIC | 16 | -117 | **-329** | -320 | -288 | TD-AL<sub>γ</sub> |
|  | 17 | -152 | -319 | **-324** | -226 | TD-AL<sub>0</sub> |
|  | 18 | -68.3 | **-225** | -198 | -140 | TD-AL<sub>γ</sub> |
|  | 21 | -92.0 | -182 | **-186** | -165 | TD-AL<sub>0</sub> |
|  | 22 | -132 | **-256** | -185 | -204 | TD-AL<sub>γ</sub> |
|  | 26 | -50.6 | **-138** | -134 | -118 | TD-AL<sub>γ</sub> |
| AIC | 16 | -125 | **-336** | -325 | -290 | TD-AL<sub>γ</sub> |
|  | 17 | -160 | -327 | **-329** | -229 | TD-AL<sub>0</sub> |
|  | 18 | -75.0 | **-231** | -202 | -142 | TD-AL<sub>γ</sub> |
|  | 21 | -98.8 | -189 | **-191** | -167 | TD-AL<sub>0</sub> |
|  | 22 | -140 | **-264** | -190 | -207 | TD-AL<sub>γ</sub> |
|  | 26 | -57.8 | **-145** | -139 | -120 | TD-AL<sub>γ</sub> |

**Table 5.** The best-fitting parameters for each mouse and model.

|  | TD-RL |  |  | TD-AL<sub>γ</sub> |  |  | TD-AL<sub>0</sub> |  | Action-Only |
| --- | --- | --- | --- | --- | --- | --- | --- | --- | --- |
| Mouse | $\kappa$ | $\alpha_v$ | $\gamma_v$ | $\kappa$ | $\alpha_h$ | $\gamma_h$ | $\kappa$ | $\alpha_h$ | $\kappa$ |
| 16 | 1.39 | 0.00297 | 1.00 | 0.0247 | 0.000479 | 0.574 | 0.0239 | 0.000190 | 0.0175 |
| 17 | 4.53 | 0.00481 | 0.869 | 0.0319 | 0.000693 | 0.143 | 0.0314 | 0.000582 | 0.0124 |
| 18 | 4.16 | 0.0227 | 0.929 | 0.0549 | 0.00274 | 0.520 | 0.0603 | 0.00164 | 0.0194 |
| 21 | 1.97 | 0.00314 | 0.927 | 0.0251 | 0.000428 | $5.71 \times 10^{-7}$ | 0.0251 | 0.000428 | 0.0153 |
| 22 | 0.992 | 0.00333 | 1.00 | 0.0250 | 0.00129 | 0.786 | 0.343 | 0.152 | 0.0132 |
| 26 | 1.94 | 0.00326 | 0.992 | 0.0399 | 0.00136 | 0.579 | 0.0354 | 0.000485 | 0.0191 |

Overall, according to both BIC and AIC measures, *all* mice preferred a TD-AL model, thus providing strong evidence that these dopaminergic dynamics represent an action-based *learning* signal, rather than an RPE or a simple representation of movement kinematics.

However, the analysis do not clearly reveal if models learning immediate or discounted action describe the dopamine activity overall better. The introduction of the third discount parameter, *γ_h_*, has an impact on model recovery. Logically, the magnitude of the best-fitting *γ_h_* parameter will likely influence the degree to which TD-AL*_γ_* is preferred relative to TD-AL_0_. So, if a large *γ_h_* value is required to replicate the dopaminergic data, the underlying model will potentially be more likely to include this term. The primary influence of *γ_h_* is the gradual transfer of PE to earlier time-points, which is visualised by the dopaminergic peak shifting to the left as training progresses. Qualitatively, mouse 22 shows the most notable forward shift in the TD-AL*_γ_* simulation, relative to the other models (Fig 5), and has the largest *γ_h_* value at 0.786, which likely account for its strong preference. It similarly has the largest difference in BIC/AIC between the two TD-AL models. It is worth noting that, for mouse 22, the best fitting TD-AL_0_ had unrealistically high *α_h_* of 0.152, where this value for all other animals was below 0.0025, which resulted in jagged curves and a lack of visual synchronicity between the simulation and the true dopaminergic data. This suggests that we failed to recover appropriate parameters for this model. Nevertheless, this mouse is well described by the TD-AL*_γ_* model.

At the other end of the spectrum, mouse 17 and 21 consistently favoured the TD-AL_0_ with both metrics. This is perhaps unsurprising given that the best-fitting *γ_h_* values were much smaller. For mouse 21, this collapsed to 5.71 ×10^-7^ , which is magnitudes smaller than the *γ_h_* values reported for all other mice. Such a small parameter value suggests that, even if TD-AL*_γ_* is the correct model used by this mouse, the difference between the outputs is statistically insignificant and so the simpler 2-parameter model will be preferred.

In sum, for all mice, we can confirm that the evidence supports the existence of an action-based learning signal which is not simply a record of current kinematics. The comparison of TD-AL*_γ_* and TD-AL_0_ is strongly influenced by the magnitude of the best-fitting *γ_h_*.

#### Group Bayesian model selection

An additional post-hoc BMS analysis was completed. Using the BIC and AIC values as a stand-in for model evidence, this method allowed the exploration of the model fits at a group level. It removes the common assumption that a single model is shared by all members of a population (a ‘fixed effects’ comparison) and, instead, calculates the likelihood of the model distribution. The protected exceedance probability, 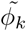, is reported in Fig 6B (see Methods for details).

**Fig 6.**
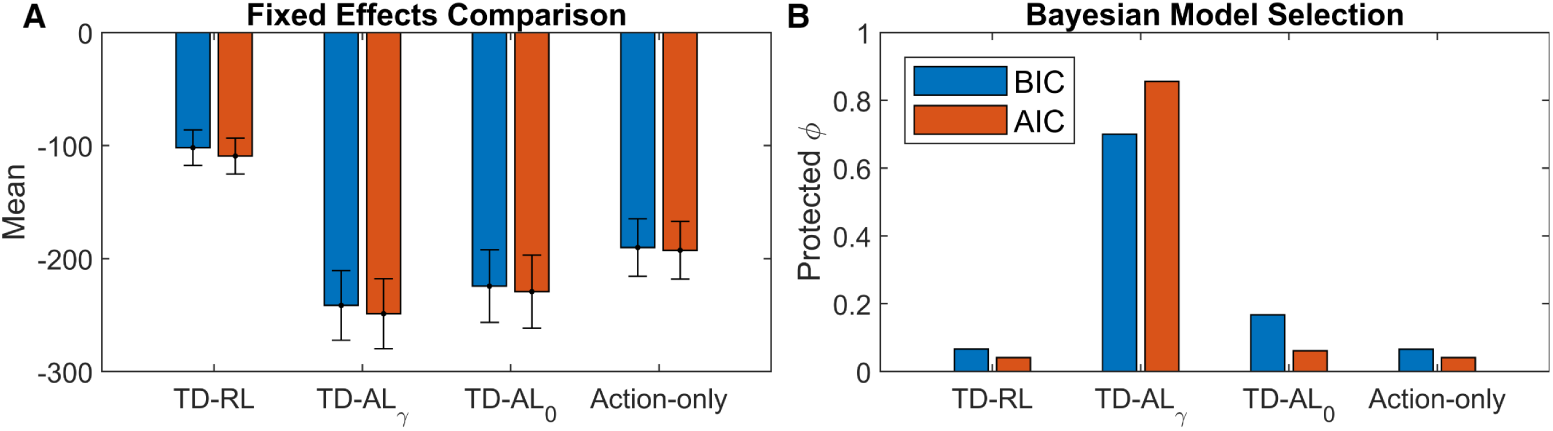
Group Analysis of Model Fits. **A:** A fixed effects comparison of the mean BIC (blue) and AIC (red) values produced by each of the models. The standard error of the mean is reported in the error bars. **B:** A comparison of the protected exceedance probabilities, 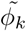, for each model using BIC and AIC as approximations for model likelihood.

For consistency with the literature, we also indicate the distribution of BIC and AIC values in Fig 6A. Under an assumption that all members of a population utilise the same model, the fixed effect comparison method [77, 78] expects the correct model to consistently have the lowest (here, most negative) BIC across all subjects.

The fixed effect analysis confirms that the non-TD-RL models have significantly lower BICs and AICs and, of these, TD-AL*_γ_* is the best fitting model on average, though this difference is negligible relative to TD-AL_0_. As expected, TD-RL performs the worst.

In contrast, the results from our BMS analysis are much clearer. Under the assumption that *some* mice may use alternative (potentially nested) models, the protected exceedance probability gives a 70.1% likelihood that TD-AL*_γ_* is the most prevalent model within the population, over and above chance. TD-AL_0_ is the next most likely at 16.7%. This separation of 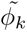 between the two TD-AL models is exacerbated when the additional parameter is less strongly penalised by AIC, with probabilities of 85.7% and 6.1%, respectively.

In all cases, TD-AL far exceeds the presence of action-only and TD-RL models, therefore providing strong evidence that, for this population of mice, the dopaminergic signal is more likely to contain an APE than an RPE.

## Discussion

Over the course of this paper, we have demonstrated how an APE-based learning signal could be applied to develop time-dependent habitual behaviours, and provided evidence for the existence of APEs within real dopaminergic data. Specifically, the novel TD-AL algorithm was tested on dopaminergic data provided by Greenstreet et al. [26]. Overall, for each of the six mice investigated, BIC analysis found that TD-AL was the preferred model over the alternatives. These analyses were particularly encouraging as the dynamics within the dopaminergic data are quite complex and TD-AL*_γ_* was able to reproduce many key features with only 3 free parameters.

### Testable model predictions

The simulations of the instrumental association task (see Results) revealed that the absence of a response to predictive cues is insufficient to disprove the existence of APEs, as this can be explained through adaptations of the discount parameter, *γ_h_*. Thus, we must interrogate alternative experiments. The current formulation of TD-AL produces several predictions that can be used to determine whether a given dopaminergic dataset contains APE signals.

#### Omission responses

TD-AL makes predictions on dopamine activity associated with omission of action or predictive cues, which are analogous to predictions of TD-RL. These predictions are illustrated Fig 7 showing signals from the trained model from Fig 2B when different elements of the trial are omitted.

**Fig 7.**
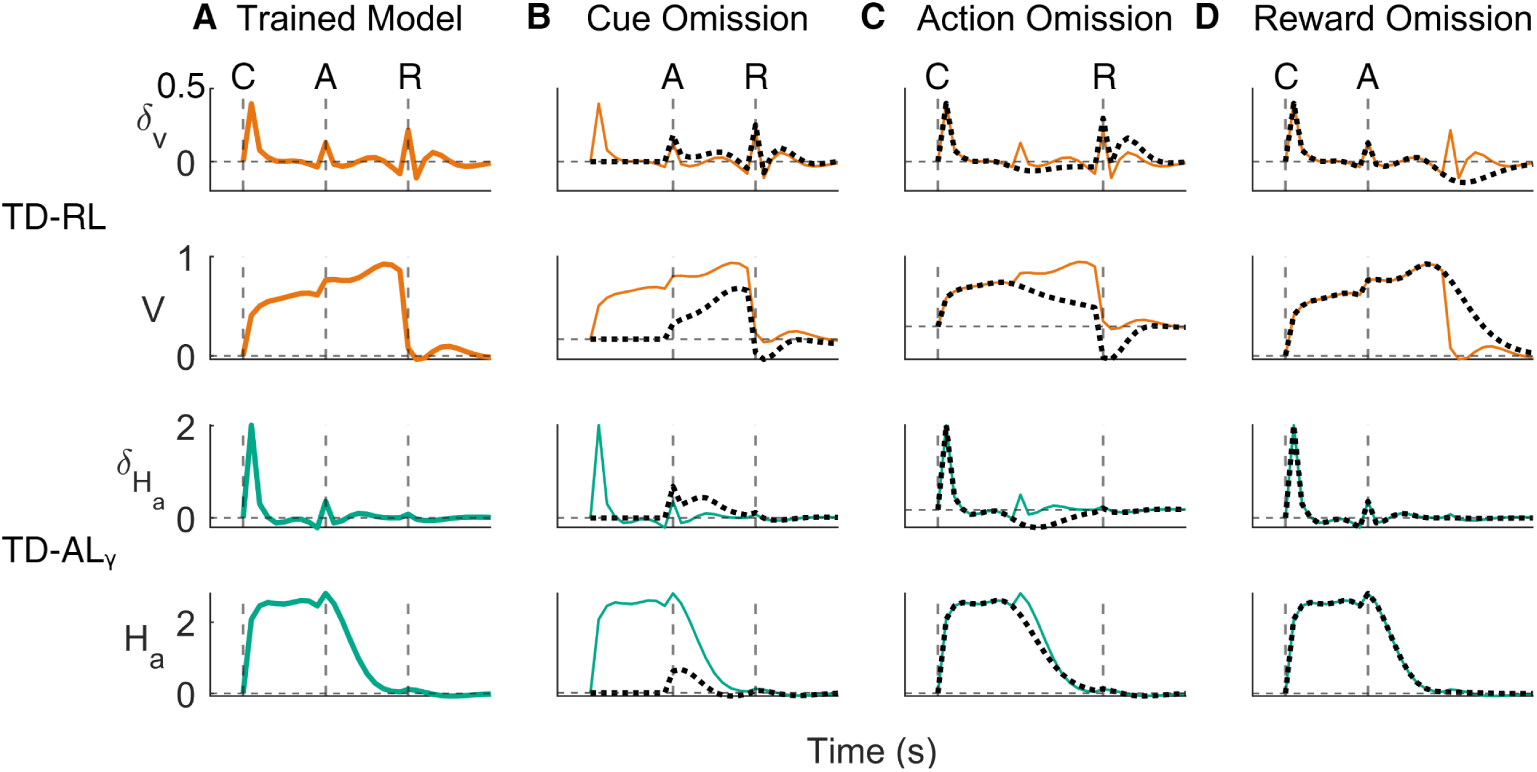
Predicted dopamine activity during omission trials. The results of simulating omission trials on the trained TD-RL and TD-AL*_γ_* in a paradigm from Fig 2. **A:** The dynamics in the final trial from the instrumental association simulations (copied from Fig 2). **B:** A single trial with cue omitted (dotted line) overlaid on the trained dynamics from A. **C:** As B, with action omitted (dotted line) rather than cue. **D:** As B, with reward omitted (dotted line).

One established property of TD-RL is that when an expected reward does not arrive, dopamine firing is partially suppressed [10] (Fig 7D). Further, the absence of associated cues will decrease the predicted future reward, and so, an RPE will increase when/if the reward arrives (Fig 7B).

TD-AL makes analogous predictions. When the predictive cue is omitted, TD-AL*_γ_* produces no response at the usual timing of the cue, because the initiation of a trial does not produce microstimuli. As a result, arrival of the action is surprising and should produce a larger APE than for a completely trained model. However, since TD-AL*_γ_* learns a continuous action, the difference in PE is mitigated and will be smaller than for a completely unexpected action to a naive model. This is indeed seen in Fig 7B, where the action-based microstimuli result in a small surprise.

When the expected action is not performed, an APE-based algorithm should behave in a comparable way to reward omission in an RPE model. As shown in Fig 7C, there is no effect on *δ_Ha_* or *H_a_* until we surpass the usual time of action. At which point, the APE is temporarily depressed until the expectation of an action fades and *H_a_* returns to baseline. This depression of *δ_Ha_* is independent of *γ_h_*, and so should always be present during action omission for a trained TD-AL agent.

Thirdly, APE-based habits are reward-insensitive, and so, there should be no change from the usual behaviour of the model. Fig 7D clearly confirms this, by showing that reward does not predict future or current action.

#### Other experimental predictions

TD-AL can be explicitly tested for using the model predictions established above. We saw that omission of an expected action will produce a negative APE and cause dopaminergic levels to dip, irrespective of *γ_h_*’s value. Unfortunately, the data used in this paper was extremely stereotyped in the final stages of the experiment, causing the number of incorrect/omission trials to be few and far between. In future, this could be resolved by the addition of a reversal task, such that the mice learn to change their response to a new mapping and execute different choices while the habitual system continues to expect the old action.

Alternatively, it should be possible to identify TD-AL in the existent data through action grouping. Specifically, if the dopaminergic signals in the later trials were classed according to the overall speed of the trial and averaged within these groups rather than over consecutive trials, then we hypothesise that the APE behaviour would differ between the slowest and fastest trials. When the animal moves the same distance over less time, the action intensity must initially spike above what was expected and produce a positive APE. In tandem, the action will also end earlier than usual, which, theoretically, would behave the same as an omission trial since the expected action is absent, causing a negative APE to follow. The reverse would be true in actions that were slower than the typical learnt movement, as the intensity remains low and continues for longer than expected.

To test which of the models, TD-AL*_γ_* or TD-AL_0_ describes dopamine activity better, one could temporally separate cue and action. TD-AL*_γ_* predicts a response to a cue while TD-AL_0_ does not. Additionally, one could vary the amount of actions (e.g. numbers of lever presses) required following a particular cue, as TD-AL*_γ_* predicts that the response would be higher for cues that need to be followed by more action.

Finally, the existence of APE-based habits would be further supported by the discovery of *H_a_*-like signals within the striatal spiny projection neurons. Just as studies have searched for value and action-value signals within the striatum [79–81], the value-free habit hypothesis supposes that action-expectation activity should be triggered during the preparation and execution of habitual behaviours. This signal should further be entirely unmodulated by outcome and reward.

#### Habitual action sequences

Realistic habitual behaviours typically combine a sequence of movements that occur over a greater length of time. Interestingly, a natural consequence of TD-AL is that the strongest S-R relationship will be between the termination of one action and the execution of the action that follows it, as has previously been proposed by Dezfouli and Balleine [23]. Just as action microstimuli will act as the closest predictor of the remaining action intensity in TD-AL and as predictive cues for subsequent rewards in TD-RL, so too can these actions be used to build expectation for those that regularly follow. Further, under the assumption of a non-zero *γ_h_*, the APE dynamics for an extremely regular series of actions will all converge to respond at the same moment in time - at the presentation of whichever cue initiates the sequence of movements. Taken to its natural conclusion, this means that when habits are completed as expected, there should be no detectable difference in the dopaminergic dynamics between a singular action plan or a series of associated muscular movements. Indeed, omission studies would be required to disentangle these two, such that a singular movement is skipped without impacting the rest of the sequence.

### Relationship to data on SOR effects

During the early research into the classification of habits as outcome-insensitive perseverative actions, experimentalists explored how the SOR could influence habit formation [25, 82]. It has since been established that resistance to extinction is more likely to arise under interval schedules over ratio [47, 83] and in variable reinforcement conditions rather than fixed [84]. Though extended training is still required to produce these habits, overtraining alone cannot account for this effect of schedule.

One leading hypothesis that was developed to explain this pattern of behaviour posits that habit formation depends on the A-O *contiguity*. When comparing fixed and variable interval schedules, DeRusso et al. [84] determined that, when action-reward correlation is held constant between conditions, the reduced A-O contiguity in the variable setting produces an increased uncertainty regarding the timing of reward. They cited this as the causal factor for resistance to extinction. DeRusso et al.’s ‘temporal uncertainty’ hypothesis was supported by Garr et al. [85], who performed a series of experiments to explore the influence of correlation, contiguity and reward density on habit expression.

The TD-AL model can be used to partially interpret SOR effects according to DeRusso’s [84] temporal action-contiguity hypothesis, though the balance of the two systems cannot be fully explained without a specific observation function. Under a free-operant task, the goal-directed system relies on previous actions to behave as predictive cues for later rewards. When there is a consistent temporal A-O contiguity, the prediction error is minimised, the same microstimulus weights are updated and ‘*Q_a_*’ will be large and precise. Accordingly, the greater the variation in the action-reward interval, the weaker the goal-directed system will be. The habit system does not suffer this disadvantage, as both variable schedules produce linear response rates - the interval between lever-presses remains constant, and so, actions are able to serve as strong predictors for themselves. However, fixed interval responses are ‘scalloped’, which will weaken the action-to-action contiguity and *H_a_* instead must increasingly rely on the previous rewards to behave as the primary predictor.

### Relationship to other computational APE models

Lee et al. [86] also incorporated a similar ‘time-dependent’ APE model to an extended version of their feature-specific model of dopaminergic heterogeneity. They likewise found that the introduction of action-specific variables was beneficial in explaining animal behaviour and established dopamine dynamics in the DLS. In particular, they explored the influence of action-sequencing within a trial. However, this model still relies on dividing a trial into separate ‘states’, i.e., learning the transitions from cue to action to reward, rather than the continuous evolution between and within these events. A novel contribution of our paper was to assess the impact of action intensity and duration on the prediction errors across continuous time.

In their original paper, Greenstreet et al. [26] also explored an APE-based ‘value-free’ algorithm, modelled on the one initially proposed by Miller et al. [16]. They demonstrated that, in trial-by-trial simulations, the impact of their experimental manipulations on animal behaviour was best explained by a ‘dual-controller’ system which transitions from value-based control to value-free with experience. Through the introduction of TD-AL, we have been able to more thoroughly exploit the rich dopaminergic data collected by Greenstreet et al. [26], which allowed us to directly examine each mouse individually and interrogate the degree to which the phasic dopaminergic changes in the TS could result in a value-free habit. This, when taken in conjunction with their work, provides significant evidence in support of APE-like signals contained within dopaminergic dynamics, as was previously proposed [24].

### Directions of future work

TD-AL does not currently describe how value and habits influence the selection of actions. Rather, it demonstrates how habits can evolve by learning to copy the choices previously made by the goal-directed system. To determine *which* action would be selected requires a Q-variable estimating action-value and a decision of how such a *Q_a_* and *H_a_* would interact.

Though they may influence each other, the specific algorithm through which actions are selected can in many ways be orthogonal to that used for learning. Rather, the most applicable algorithm is contingent on the behaviour it aims to replicate. One core advantage of TD-AL over other APE-based algorithms is its time-continuous nature. As such, it is likely that future applications of this model will be interested in replicating within-trial behaviours, be that reaction times, action intensities or other. Any model which hopes to capitalise on that must also determine *when* this action would be initiated. For example, if an experiment measures reaction times as its dependent variable, an evidence accumulation model (EAM) will be more appropriate than the softmax choice function. Thus, TD-AL lends itself to the RL-EAM family of observation functions [87], with a *H_a_* dependent drift-rate working in tandem with a corresponding *V* or *Q_a_* value. It would be particularly interesting in future research to explore how the influence of predictive cues on the temporal dynamics of *H_a_* versus *V* could influence error rates and reaction times using these more mechanistic models of action selection.

## Conclusion

This paper has extended the computational models of value-free habit formation to describe learning within a trial and predict temporal profile of an action prediction error. This model was then shown to capture dopamine activity in the tails of striatum more accurately than the equivalent value-based model. These results provide a strong proof-of-concept that habitual behaviours can be explained through value-free learning systems and that the discrepancy between the experimental and computational definition of habits can be solved through the introduction of action prediction errors.

## Materials and methods

### Fitting procedure

The dataset provided by Greenstreet et al. [26] is rich, with multiple mice completing roughly 10,000 trials while camera tracking and dLight measurements were recorded. This section outlines the key details of the additional data preprocessing completed and fitting procedure that was applied to match model prediction errors to the dopaminergic signals. A schematic overview of the fitting procedure is provided by Fig 8.

**Fig 8.**
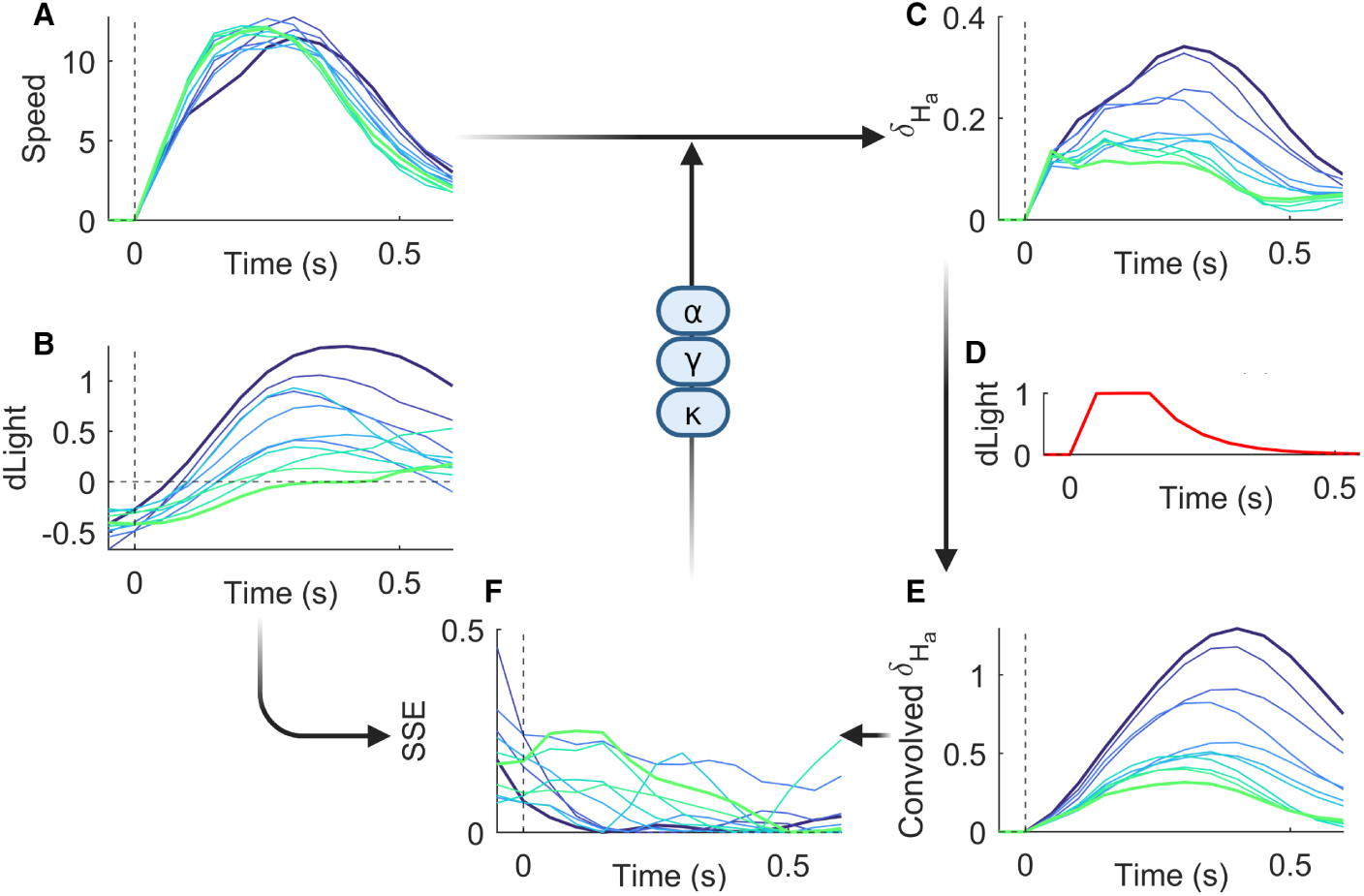
Model Fitting Procedure. **A:** Camera tracking data was used to extract the speed of the mouse (averaged between nose and ears) from the time it left the centre port (0s) until it entered the side port. **B:** The dLight data processed as in Fig 3B. **C:** For a given set of parameters, the model learns from the behavioural data and the associated PE is calculated. These trials are then averaged over 200 consecutive events. **D:** The temporal dynamics of dLight fluorescence. This is convolved with the PE to produce an equivalent signal to the experimental data, as though the same methodology was used to record both the PE and dopaminergic data. **E:** The resulting convolved PE. **F:** The cost function is calculated as the sum squared error (SSE) of the averaged dopaminergic signals (**B**) and convolved PEs (**E**) in the 0.5s following action initiation. The parameters update in order to minimise this cost. **C-F** repeats until the best-fitting parameters are returned.

#### Behavioural simulation

Each learning model was provided with a simplified series of continuous events determined by the real experiences of a given mouse. Thus, as illustrated in Fig 9, trials obeyed the following sequence of steps:

1. An auditory cue sounds (either high or low).
2. The mouse exits the centre nose port (the action is initiated) and the choice is determined by the port entered (either left or right).
3. The continuous intensity of the trial’s action is set to the mouse’s speed, as extracted from the camera tracking data and averaged between the nose and ear movement.
4. If the mouse makes the correct action, it receives a reward.

**Fig 9.**
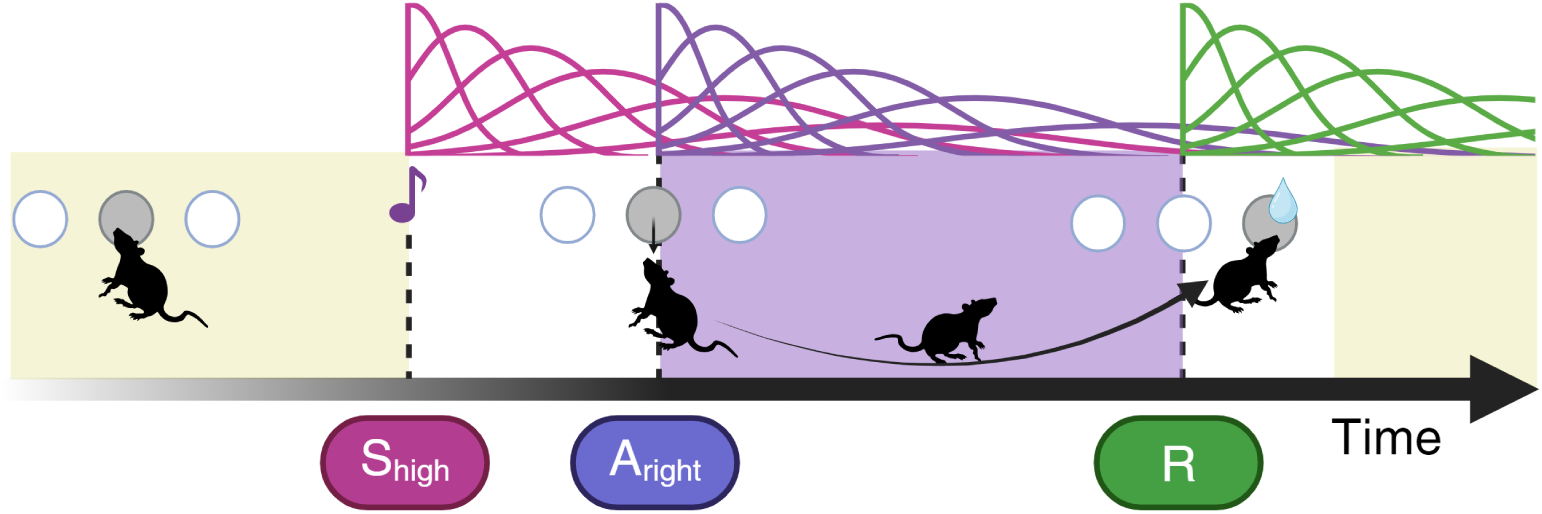
A Simulated Trial. Following an ITI (yellow), during which all previous microstimuli are allowed to finish and the next trial begins, the ‘go’ tone initiates the cue microstimuli (associated with either *S*_high_ or *S*_low_). The action microstimuli (*A*_left_ or *A*_right_) begin when the mouse leaves the centre port. The action intensity dynamics, *A_a_*(*t*), are determined by the speed of the mouse and continue until the mouse enters the associated port. If the correct choice is made, a reward (*R*) is received, the reward microstimuli are produced and the next ITI begins.

In total, there were five potential events that produced microstimuli (two cues, two actions and a reward), resulting in 60 weights, *w_ij_*, (*m* = 12 per event) which update at every timestep (dt = 0.05s) according to either Eq 7 or Eq 9. The delays between these events were determined by those actually experienced by the mouse on the associated trial. At the end of the trial, a fixed ITI of 30dt (1.5s) was given, such that all microstimuli from the previous trial had ended and the model could not learn to expect the subsequent auditory cue.

While training the model, action intensity *A_a_*(*t*) in each time step on each trial was set to animals speed. This value was averaged between a mouse’s ears and computed from the camera tracking data simply as the Euclidean distance a body point travelled in each frame, multiplied by the frame rate. Speed was selected as the measure of action intensity for two reasons. First, speed is a continuous measure which remains positive regardless of the choice made, whereas position, for example, would need to be corrected relative to a starting point. This allowed for minimal preprocessing to be required and for the two actions’ *H_a_* variables to receive equivalent information. Second, experience and training lead to an increased speed and reduced trial duration [26]. In using this measure directly as the learnt output, we can partially control for these training effects on the APEs and resulting dopaminergic signals.

#### Processing dLight data

All preprocessing described by Greenstreet et al. [26] had been completed prior to our data access. The original dopaminergic signal had a resolution of 10,000 Hz and so, following a rolling Z-score across a window of 1.6s, the frequency was reduced to a timestep of 0.05s via MATLAB’s `resample` function, which uses linear interpolation to calculate new values based on averages from the true datapoints.

To reproduce Greenstreet et al.’s [26] results for each mouse (as in Fig 3B) the dopaminergic data was divided into individual trials comprising of a 1.5s window centred around action initiation. Using the subset of trials in which the contralateral port was cued, the dopaminergic curves were averaged across 200 consecutive trials to produce a series of movement-locked peaks. A total of six mice completed this task with TS recordings, and their individual datasets are visualised in Fig 5.

#### Fitting the prediction errors

Once a simulation is run with the given dataset and model parameters (*α* and *γ*), the resulting PE signal can be compared to the dopaminergic data to calculate the goodness-of-fit.

The time course of the dLight fluorescence contaminates the dopaminergic signal with a consistent artifact. Thus, to produce comparable PE signals, we convolved the *δ* variable with a kernel representing the temporal effect of dopamine binding on the sensor (Fig 8D), using MATLAB’s `conv` function. The rise and fall time of these kernels were set to 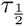 = 9.5ms and 90ms, respectively, as reported in Patriarchi et al. [70]. The convolved PE signal is processed in an equivalent manner to the dopaminergic signal to produce trial averages over training, as shown in Fig 5. The cost function of the current simulation is calculated as the SSE in the 0.5s (10 timesteps) following action initiation for the averaged dopaminergic data and the averaged, convolved PE data, as specified in Eq 12 (and mentioned in Fig 8E):

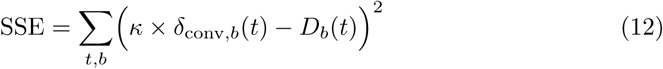

where: SSE = cost function for a given dataset and parameter combination,

*δ*_conv*,b*_ = convolved PE signal, averaged over trials in block *b*,

*D_b_* = dopaminergic signal, averaged over trials in block *b*,

*t* = timestep,

*b* = block number (each block consists of 200 trials)

*κ* = scaling factor.

Note that the prediction error is multiplied by a scaling parameter, *κ*, to minimise the difference between the two figures. Its value can be analytically determined, via Eq 13.

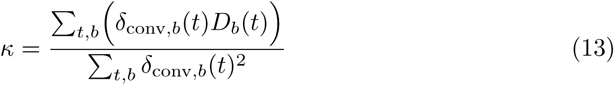

This process can be conceptualised for a TD-AL learning model using Algorithm 1 (the signals entering the cost function are computed based on blocks of 200 trials, and final trials beyond the last block of 200 trials are excluded from fitting).

The cost function is minimised over several iterations of the above simulations from 20 randomised start values, using MATLAB’s fmincon function, until the best-fitting parameters are established.

### Statistical methods

#### Individual Mice

The recovered model for each mouse is determined via BIC analysis [88], such that a lower BIC implies a better fitting model, though the AIC [89] is also reported. These values represent a measure of model fit that includes a penalty for additional parameters to counteract the risk of overfitting. As our likelihood measure is approximated using

#### Algorithm 1

**Evaluating how well specific model parameters fit data from one mouse.**

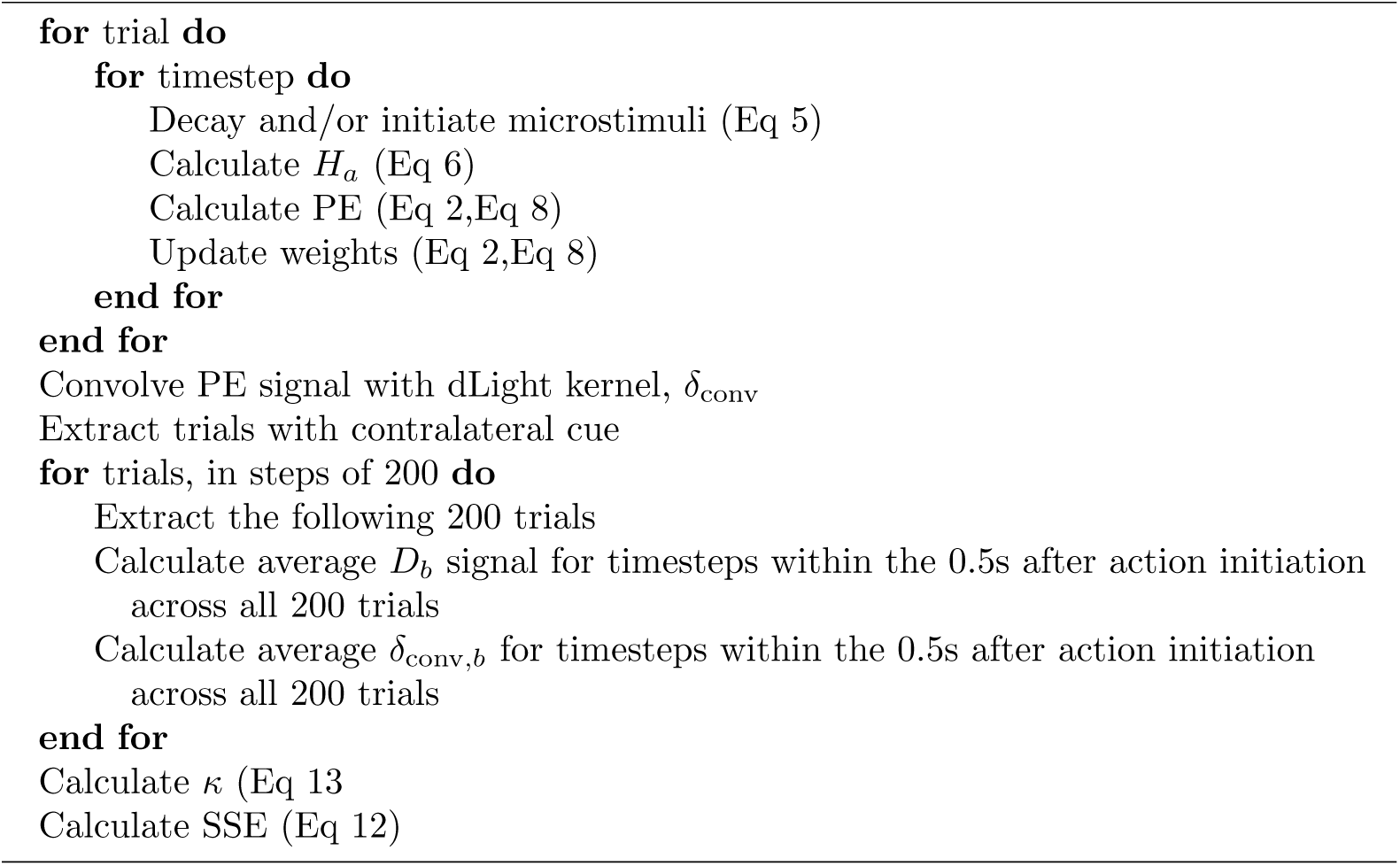

an SSE cost function, the equations to calculate BIC and AIC differ slightly from the standard in order to account for the Gaussian distribution of errors [90]. Thus, the values reported in this chapter are calculated using Eq 14 and Eq 15:

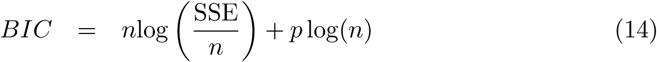

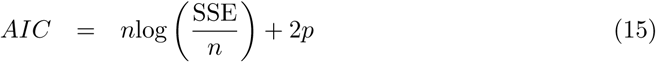

where: *p* = number of free parameters in the model,

*n* = number of datapoints used to calculate the SSE.

It is worth noting that, when the number of datapoints used to calculate BIC is large, BIC may over-penalise additional parameters and incorrectly prefer a model that is too simple, though this is partially mitigated by our use of the SSE-adapted BIC calculation (Eq 14). This is a particular risk here, as we use many datapoints per trial and so the BIC values are penalised twice as strongly as AIC. Further, given that the dopaminergic data is quite complex and the maximum number of parameters for any model is three, overfitting due to increased model complexity is not likely. Thus, we also report the AIC values in Table 4 and consider these results when looking at the visualisation of the best-fitting model simulations.

#### Measuring recovery

The above fitting procedure was assessed through *parameter* and *model* recovery analyses, in which surrogate data is produced with a known model and parameter values. Parameter estimates and BIC values can be extracted for each of the simulated datasets by applying the fitting procedure as though we were working with real behavioural data.

Due to the absence of an ‘action-selection’ algorithm for this experiment through which a mouse’s behaviour could be simulated, we instead produced surrogate ‘dLight’ signals for each mouse using the behavioural simulation method outlined above. An agent applies a known learning algorithm with preset parameter values to the real behavioural data. The resulting PE is convolved with a dLight kernel and then treated as though it was the true dopaminergic signal. For each set of parameter values, each model is independently simulated to create 6 individual datasets based on the behaviour of each mouse (a dataset consists of between 3700-5500 trials). Each of the four models is then separately fitted to every individual dataset, using the same fitting procedure that is used for real data (Algorithm 1). The parameter values used to create the surrogate datasets are outlined in Table 8.

**Table 8.** Parameter combinations used to simulate prediction errors produced from Greenstreet et al. [26] data.

| True Model | True $\alpha$ | True $\gamma$ | Datasets per mouse |
| --- | --- | --- | --- |
| TD-RL | 0.025,0.05,0.5,0.75, 1 | 0, 0.25, 0.5, 0.75, 0.97 | 25 |
| TD-AL$_{\gamma}$ | 0.025,0.05,0.5,0.75, 1 | 0, 0.25, 0.5, 0.75, 0.97 | 25 |
| TD-AL$_0$ | 0.025,0.05,0.5,0.75, 1 | 0 | 5 |
| Action-only | 0 | 0 | 1 |

The most right column shows the total number of simulated datasets per mouse. For the models with two parameters, each potential combination was utilised. The difference in parameter-space dimensions means that fewer surrogate datasets were created for TD-AL $_0$ and the action-only model.

All four models are then tested on this surrogate data using the fitting procedure, which resulted in a total of 1344 sets of estimated parameters and BIC values. Using these results, parameter recovery is straightforward to assess from the subset of agents using the same model as the data - the correlation between true and estimated parameter values should approach a perfect positive correlation as the accuracy increases. Model recovery is usually quantified in two ways, through a *confusion matrix* and/or an *inverse matrix* [78]. For each surrogate dataset, BIC analysis is used to select the best-fitting model. By comparing the true model, *M_T_*, with the recovered model, *M_R_*, we can calculate both the *P* (*M_R_|M_T_*) and the *P* (*M_T_|M_R_*), for every model. The former of these is the most commonly reported in a confusion matrix, while the latter inverts the confusion matrix to provide a measure of our confidence that the recovered model truly is underlying the data. As the accuracy and reliability of model selection increases, the mean value of the matrices’ diagonals will also increase, thereby allowing us to compare a single measure when discussing recovery efficiency.

#### Group Bayesian Model Selection

The BMS analysis was used to compare model likelihood at a group level (see Stephan et al. [77] and Rigoux et al. [68] for full details). This method removes the common assumption that a single model is shared by all members of a population (a ‘fixed effects’ comparison) and, instead, calculates the likelihood of the model distribution.

BMS assumes that the ratio of models within a population takes the form of a Dirichlet distribution, such that each model, *k*, occurs with a probability, *r*, given the ‘concentration’ of that model in the population, *α_k_* (Eq 16). Additionally, this method presumes that the models compared are the only models present in the population and thus, all *r* sum to 1.

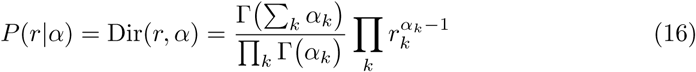

where:

*r* = probability of model in population,

Γ = the gamma function,

*α* = unobserved occurrence of models in population,

*k* = a given model within the population set.

Stephen et al. [77] proposed that the model likelihoods can be compared according to the conditional model probability, *P* (*r*|*y*; *α*), where *y* represents the actual data observed. The resulting exceedance probability, *ϕ_k_*, can be interpreted as a measure of how likely it is that a given model, *k*, is the most prevalent model in a population (Eq 17).

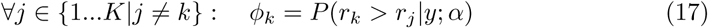

where: *ϕ_k_*= exceedance probability for the model, *k*,

*K* = total number of models.

Rigoux et al. [68] further extended this measure to produce the *protected* exceedance probability, 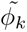, which considers an error rate, or ‘Bayesian omnibus risk’, and calculates how likely a given model, *k*, is to be the most prevalent model, *over and above chance* (Eq 18). This can be conceptualised as an alternative to calculating a p-value.

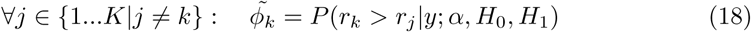

where:

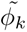 = protected exceedance probability for the model, *k*,

*H*_0_ = null hypothesis that all models are equally frequent,

*H*_1_ = alternative hypothesis that all models are not equally frequent.

These were calculated using the bms function created by Gershman [91].

## Acknowledgments

The authors would like to acknowledge the use of the University of Oxford Advanced Research Computing (ARC) facility in carrying out this work. (https://doi.org/10.5281/zenodo.22558).

